# XPO1 controls VDR localization and activity in triple-negative breast cancer cells

**DOI:** 10.64898/2026.10.07.756946

**Authors:** J Denis, S Bonnet, B Győrffy, U Jeschke, C Teyssier, V Cavaillès, S Jalaguier

## Abstract

The association of cytoplasmic localization of VDR in TNBC tumors and disease aggressivity prompted investigation of its regulation by XPO1. In TNBC cell lines, the XPO1 inhibitors selinexor and eltanexor increased nuclear VDR localization. Their effects were particularly strong when combined with vitamin D treatment. Similarly, XPO1 knockdown increased nuclear VDR levels, both alone and in combination with vitamin D. These results identify XPO1 as an important regulator of VDR subcellular distribution. XPO1 inhibition also enhanced vitamin D-dependent VDR transcriptional activity. Indeed, selinexor, eltanexor, and XPO1 silencing increased vitamin D-dependent CYP24A1 reporter activity and the expression of genes such as *CYP24A1* or *CDH1*. RNA sequencing showed that XPO1 modulation markedly reshaped the vitamin D transcriptional response. The affected genes were mainly involved in metabolism, cell-cycle regulation, protein localization, and cellular organization. Functional analyses demonstrated that cytoplasmic VDR promoted TNBC cell proliferation. Conversely, nuclear VDR was required for the antiproliferative effects of vitamin D. Accordingly, XPO1 inhibition or depletion potentiated vitamin D-mediated growth suppression. This finding suggested functional cooperation between XPO1-dependent export and VDR activity. Finally, high VDR expression was associated with poor disease-free or overall survival only in tumors with high XPO1 expression. Overall, XPO1 negatively regulates VDR nuclear localization and vitamin D responsiveness in TNBC. Targeting XPO1 may therefore enhance the therapeutic potential of vitamin D-based strategies in TNBC.

## INTRODUCTION

Breast cancer represents approximately 25 % of newly diagnosed female cancer worldwide ^1^. It is a highly heterogeneous disease with subtypes classified according the expression of estrogen receptor (ER), progesterone receptor (PR), human epidermal growth factor receptor 2 (HER2) and the proliferation marker Ki67. Among these subtype, 15-20 % are triple negative breast cancers (TNBC; ER-, PR-, HER2-) ^2^. The absence of expression of the three receptors renders conventional hormone therapies as well as HER2-targeted therapies ineffective ^3^. As a result, TNBC patients who are not eligible for surgery undergo nonspecific chemotherapy. However, less than 30 % of patients with TNBC achieve a complete response and both recurrence and mortality rates are significantly higher than those of other breast cancer subtypes ^4^. Although inhibitors of bromodomain and extra-terminal domain (BET) family of proteins such as iBET151 or OTX015proved their efficacy in some TNBC cases, rapid resistance to these drugs frequently develops through diverse mechanisms ^5^. Adverse features such as drug resistance, high heterogeneity and malignancy highlight the need for combinatorial targeted approaches to improve TNBC outcomes ^4^.

Numerous epidemiological studies have established an inverse association between serum vitamin D and breast cancer risk. Moreover, vitamin D deficiency is a common feature of TNBC patients ^6^. The active form of vitamin D, 1,25(OH)2D3 exerts its effects through binding to the vitamin D receptor (VDR) which subsequently forms a heterodimer with the retinoid X receptor (RXR). Notably, heterodimerization between VDR and RXR is not permissive and the complex cannot be activated by the RXR ligand, 9-*cis* retinoic acid ^7^.

Vitamin D has been shown to exert anti-proliferative effect on breast cancer cells *in vitro* ^8^ and *in vivo* as demonstrated in different transgenic mouse models ^9^. VDR also regulates apoptotic pathways by downregulating expression of anti-apoptotic factors such as Bcl-2 and/or increasing expression of pro-apoptotic genes including Bax or Bak ^10^. In breast cancer tissue, VDR expression inversely correlate with cancer aggressiveness with benign breast lesions displaying a stronger expression than *in situ* or invasive carcinomas. Additionally, VDR expression is higher in luminal A breast cancer than in TNBC ^11^.

In a recent study, we showed that cytoplasmic VDR expression in TNBC patients is significantly associated with reduced disease-free survival ^12^. In ER-positive BC cytoplasmic localization of VDR is linked to aggressive disease progression whereas nuclear VDR appears to have a protective role ^12^. Nucleocytoplasmic trafficking is mediated by various members of the Karyopherin-β (Kap) family of nuclear transport receptors, which includes importins, biportins and exportins ^13^. Among the exportins, XPO1, also known as chromosomal region maintenance 1 (CRM1), is a key nuclear export protein that recognizes nuclear export signals present in numerous cargoes, including tumor suppressors and cell cycle regulators, as well as RNAs and mediates their transport them to the cytoplasm in a Ran GTPase-dependent manner ^14,15^. Elevated XPO1 expression has been associated with poor survival and increased risk of recurrence in tamoxifen-treated breast cancer patients ^16^.

Selinexor (KPT-330), a first-in-class selective inhibitor of nuclear export (SINE) targeting XPO1, received FDA approval in 2020 for the treatment of multiple myeloma ^17^. Notably, selinexor significantly reduces the viability of various TNBC cell lines, in part through inhibition of the NF-κB pathway ^18^. A second-generation SINE, eltanexor (KPT-8602), has recently been evaluated in patients with hypomethylating agent-refractory higher-risk myelodysplastic syndromes and was well tolerated ^19^.

In the present study, we show that, in TNBC, high VDR expression is associated with poor prognosis in tumors with elevated XPO1 expression suggesting that the regulation of VDR subcellular localization by XPO1 might be critical for its role in BC progression. Using MDA-MB-231 and MDA-MB-436 TNBC cells, we further demonstrate that SINE compounds, as well as XPO1 knockdown, significantly enhance vitamin D-dependent nuclear localization of VDR. As a result, ligand-dependent VDR transcriptional activity is markedly increased following treatment with selinexor, eltanexor, or siXPO1. A transcriptomic analysis revealed that pathways such as metabolic process or cell cycle are affected by vitamin D upon downregulation of XPO1. Accordingly, inhibition or downregulation of XPO1 amplifies vitamin D-mediated suppression of TNBC cell proliferation. This work opens up new perspectives in the fight against TNBC by proposing a novel easily druggable therapeutic target.

## MATERIALS AND METHODS

### Materials

1α,25-dihydroxyvitamin D3, selinexor and eltanexor were purchased from Sigma-Aldrich, Euromedex and Selleckchem respectively.

### Cell culture and transfections

MDA-MB-436 (RRID:CVCL_0623) and MDA-MB-231 (RRID:CVCL_0062) were cultured in RPMI (Invitrogen) supplemented with 10 % fetal calf serum (FCS) and antibiotics (Gibco). All cell lines were authenticated by short tandem repeat profiling and free from mycoplasma. When needed, plasmids or siRNAs were transfected using either JetOPTIMUS or INTERFERin (Polyplus), respectively. When indicated, cells were treated with vitamin D (Sigma-Aldrich) and/or selinexor or eltanexor (Sellekchem) and then harvested 24 hours later. The list and sequences of the siRNAs used in this study are indicated in Supplementary table 1. SiCtl is control siRNA, designed not to target any gene.

### Immunofluorescence

MDA-MB-436 and MDA-MB-231 cells were either transfected with siXPO1 or siCtl and then treated with vitamin D or cotreated with selinexor or eltanexor and vitamin D. IF revelation was then performed using an anti-VDR antibody (rabbit monoclonal antibody D2K6W) (Cell Signaling Technology). The cells were also stained with Hoechst (Sigma-Aldrich) and mounted for fluorescence microscopy using a Zeiss Axioimager M2 Apotome2. Staining quantification was done using the Fiji software.

### mRNA quantification

RNA was isolated using the Zymo Research kit (Zymo research) and reverse transcription (RT)-qPCR assays were done using OneScript (Ozyme) according to the manufacturer’s protocol. Transcripts were quantified using SensiFAST SYBR (Ozyme) on a LC480 instrument. The nucleotide sequences of the primers used for real time PCR are indicated in Supplementary Table 1.

### Luciferase assays

MDA-MB-436 and MDA-MB-231 cells were plated in 96-well plates 24 hours prior to transfection with either JetOPTIMUS® (Polyplus) or INTERFERin® (Polyplus) according to the manufacturer’s protocol. Firefly luciferase data were normalized to renilla luciferase activity and expressed as relative luciferase activities.

### Cell proliferation assay

MDA-MB-436 and MDA-MB-231 were seeded at a density of 2000 cells per well. At the indicated time, AlamarBlue (Invitrogen^TM^) was added to selected wells and cells were incubated at 37°C for 1h. Fluorescence is measured using excitation at 560 nm and emission at 590 nm on a spectrophotometer. Results were normalized to the cell density at day 1.

### RNA sequencing and data analysis

Thirty total RNA samples from MDA-MB-436 breast cancer cells were analyzed, representing ten experimental conditions with three biological replicates each: control, selinexor, eltanexor, vitamin D, their respective combinations with vitamin D, siCtl, siCtl + vitamin D, siXPO1, and siXPO1 + vitamin D.

RNA quality was assessed using Qubit® 2.0. Libraries were prepared and sequenced by Helixio (Clermont-Ferrand, France) on an Element AVITI™ instrument using paired-end 2 × 75-bp sequencing. Reads were quality-controlled with FastQC and MultiQC, trimmed with Trimmomatic, and aligned to the human GRCh38 reference genome (release 115) using STAR. Reads shorter than 50 bp after trimming were discarded. Gene-level counts were generated with HTSeq-count in union mode, using stranded counting and a minimum alignment quality of 10; multimapping and ambiguous reads were excluded.

Count matrices were analyzed with DESeq2 after filtering low-expression genes. Variance-stabilized data were used for principal component analysis and hierarchical clustering. Differential expression was assessed using DESeq2 normalization and pairwise comparisons evaluating XPO1 modulation, vitamin D treatment, and their combination. Genes with an adjusted p<0.05p < 0.05p<0.05 and ∣log 2FC∣>1|\log_2\mathrm{FC}| > 1∣log2FC∣>1 were considered differentially expressed.

Gene Ontology Biological Process enrichment was performed separately for upregulated and downregulated genes using topGO in Galaxy, with Fisher’s exact test and Benjamini–Hochberg correction. Significantly enriched terms were defined by an adjusted p<0.05p < 0.05p<0.05 and visualized as dot plots using SRplot.

### Survival analysis

Using the KM-plot Private Edition (http://kmplot.com), Affymetrix and RNA sequencing data from the Cancer Genome Atlas (TCGA) were analyzed using Cox proportional hazards regression, as previously described (ref). The Kaplan-Meier method was used to estimate overall survival calculated from diagnosis to death. Patients lost to follow-up were censored at the time of last contact.

### Statistical analysis

All experiments were conducted independently at least three times. Results were expressed as the mean ± standard deviation (S.D.). Comparisons of two independent groups were performed using the Mann-Whitney U test. A probability level (*p* value) of 0.05 was chosen for statistical significance. Statistical analyses were performed using StatEL (AdScience).

## RESULTS

### XPO1 inhibition increases vitamin D-dependent nuclear localization of VDR

The observation of cytoplasmic expression of VDR in TNBC being associated to a more aggressive disease ^12^prompted us to investigate the regulation of VDR subcellular localization. Exportins are well-established regulators of nucleocytoplasmic trafficking, and XPO1 expression has been reported to be adversely associated with breast cancer ^16^.

To investigate the ability of XPO1 to regulate VDR subcellular localization, we first assessed the effect of two selective inhibitors of nuclear export (SINE), selinexor and eltanexor. Two TNBC cell lines, MDA-MB-436 (Figure 2A) and MDA-MB-231 (Figure 2B), were treated with these inhibitors and subjected to immunofluorescence staining using an anti-VDR antibody. Representative images showing VDR and Hoechst staining, along with merged images, are displayed on the left panels, while quantifications are presented on the right panels.

**Figure 1.**
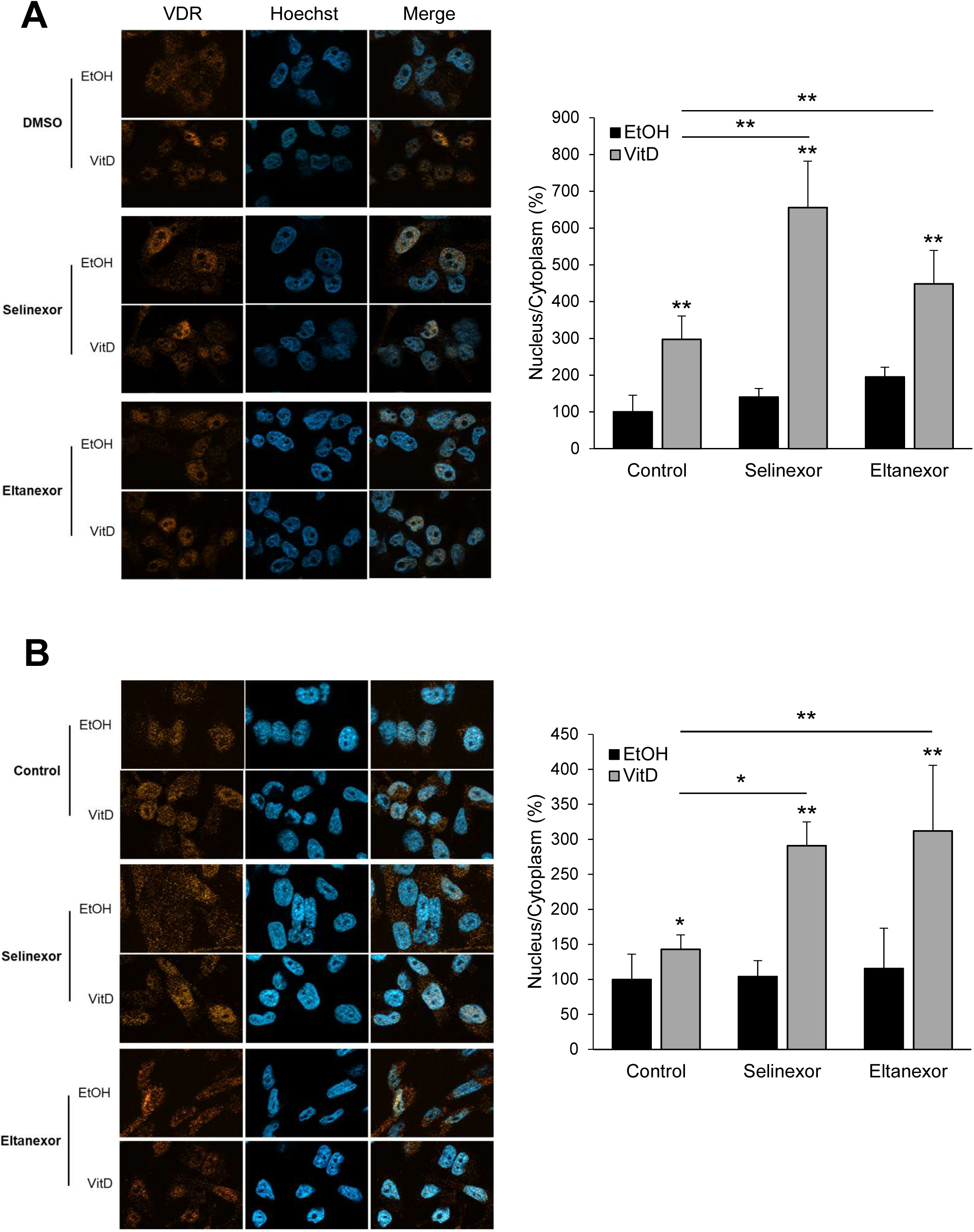
selinexor and eltanexor augment VDR nuclear localization. MDA-MB436 (A) and MDA-MB-231 (B) were treated with either selinexor (10^-8^ M) or eltanexor (10^-8^ M) and/or vitamin D (10^-9^ M). Immunofluorescence staining of MDA-MB-436 and MDA-MB-231 cells (left panels) was performed with an anti-VDR antibody and nuclei were counterstained using Hoechst. Cells were fixed after 24 h of treatment, and images were acquired at 40× magnification using a Zeiss AxioImager M2 Apotome2. Staining quantification (right panels) was performed using Fiji software, and results are expressed as a percentage relative to the ethanol (EtOH)-treated control condition. Statistical analyses were performed using the Mann-Whitney test. *p<0.05; **p<0.01.

**Figure 2.**
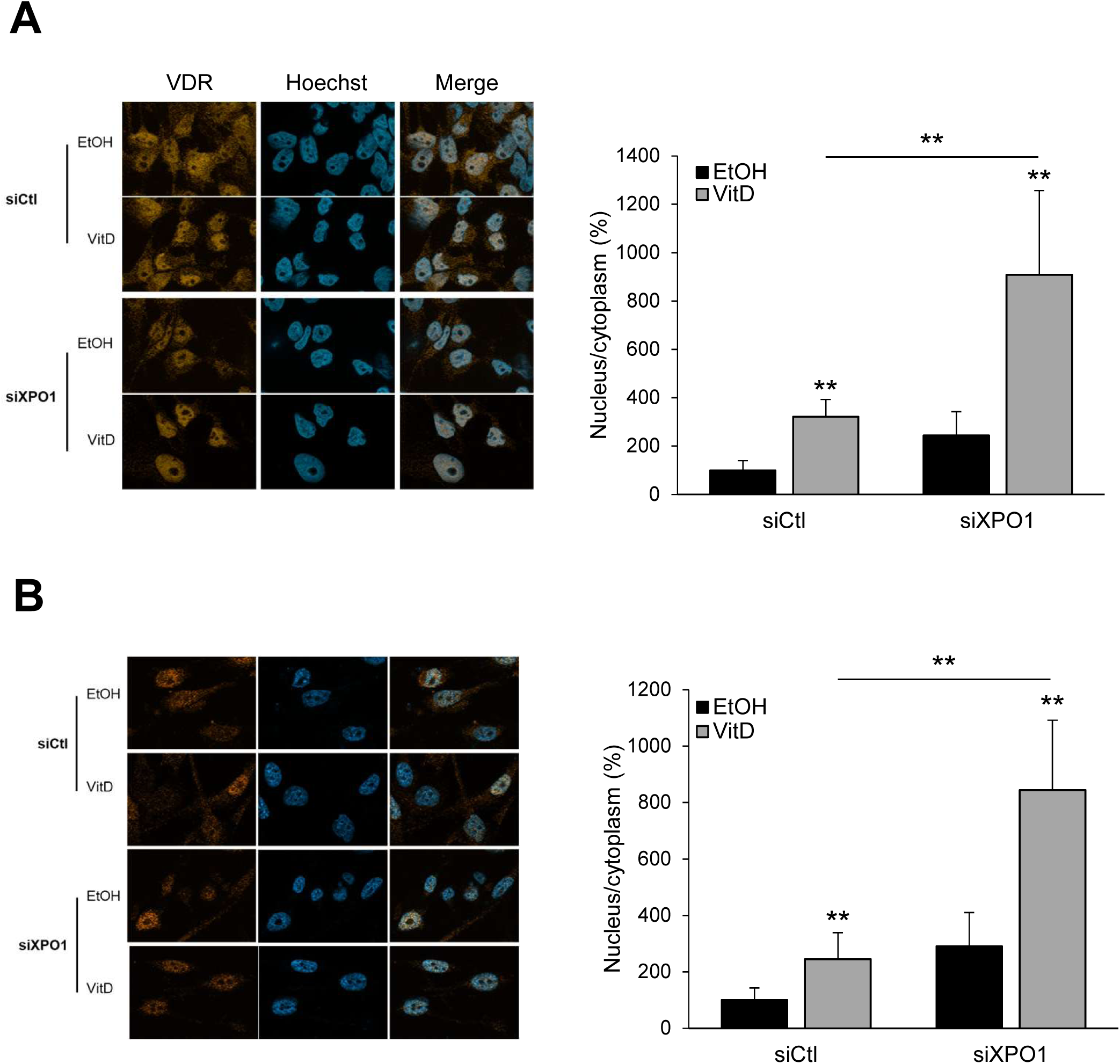
Down-regulation of XPO1 expression increase VDR nuclear localization. MDA-MB436 (A) and MDA-MB-231 (B) were transfected with siRNAs, control (siCtl) or targeting XPO1 (siXPO1) and treated or not with vitamin D (10^-9^ M). Immunofluorescence staining and quantification were done as in Figure 2. Statistical analyses were performed using the Mann-Whitney test. **p<0.01.

In MDA-MB-436 cells (Figure 2A, right panel), vitamin D treatment increased nuclear VDR localization by approximately threefold compared to ethanol control. Selinexor alone had a modest effect (1.4-fold), whereas the combination of vitamin D and selinexor showed a strong synergistic effect (6.6-fold increase). Eltanexor alone increased nuclear VDR levels by nearly twofold, and its combination with vitamin D further enhanced this effect compared to vitamin D alone. In MDA-MB-231 cells (Figure 2B, right panel), vitamin D alone had a modest effect (1.4-fold). Neither SINE alone significantly affected nuclear VDR localization. However, combining vitamin D with either selinexor or eltanexor resulted in a clear synergistic increase. In MCF7 cells, vitamin D alone increased nuclear VDR (1.8-fold), selinexor had a modest effect, and the combination further enhanced nuclear localization (2.7-fold compared to vitamin D alone) (Supplementary Figure 1A). These results indicate that pharmacological inhibition of XPO1 activity enhances nuclear VDR expression.

### XPO1 knock-down augments ligand-dependent nuclear localization of VDR

We then examined whether genetic downregulation of XPO1 would produce similar effects. Knockdown efficiency was confirmed (Supplementary Figure 1B), showing approximately 60% reduction at both mRNA and protein levels. MDA-MB-436 (Figure 3A) and MDA-MB-231 (Figure 3B) cells were transfected with siXPO1 and treated with vitamin D. In control siRNA conditions, vitamin D increased nuclear VDR levels (3.2-fold in MDA-MB-436 and 2.5-fold in MDA-MB-231), and siXPO1 alone also increased nuclear VDR (2.4-fold and 2.9-fold, respectively). The combination of siXPO1 and vitamin D resulted in a strong synergistic effect (9.1-fold and 8.4-fold increases). Similar effects were observed in MDA-MB-468 cells (Supplementary Figure 1C). Together, these results demonstrate that XPO1 is a key regulator of VDR subcellular localization in TNBC cells.

**Figure 3.**
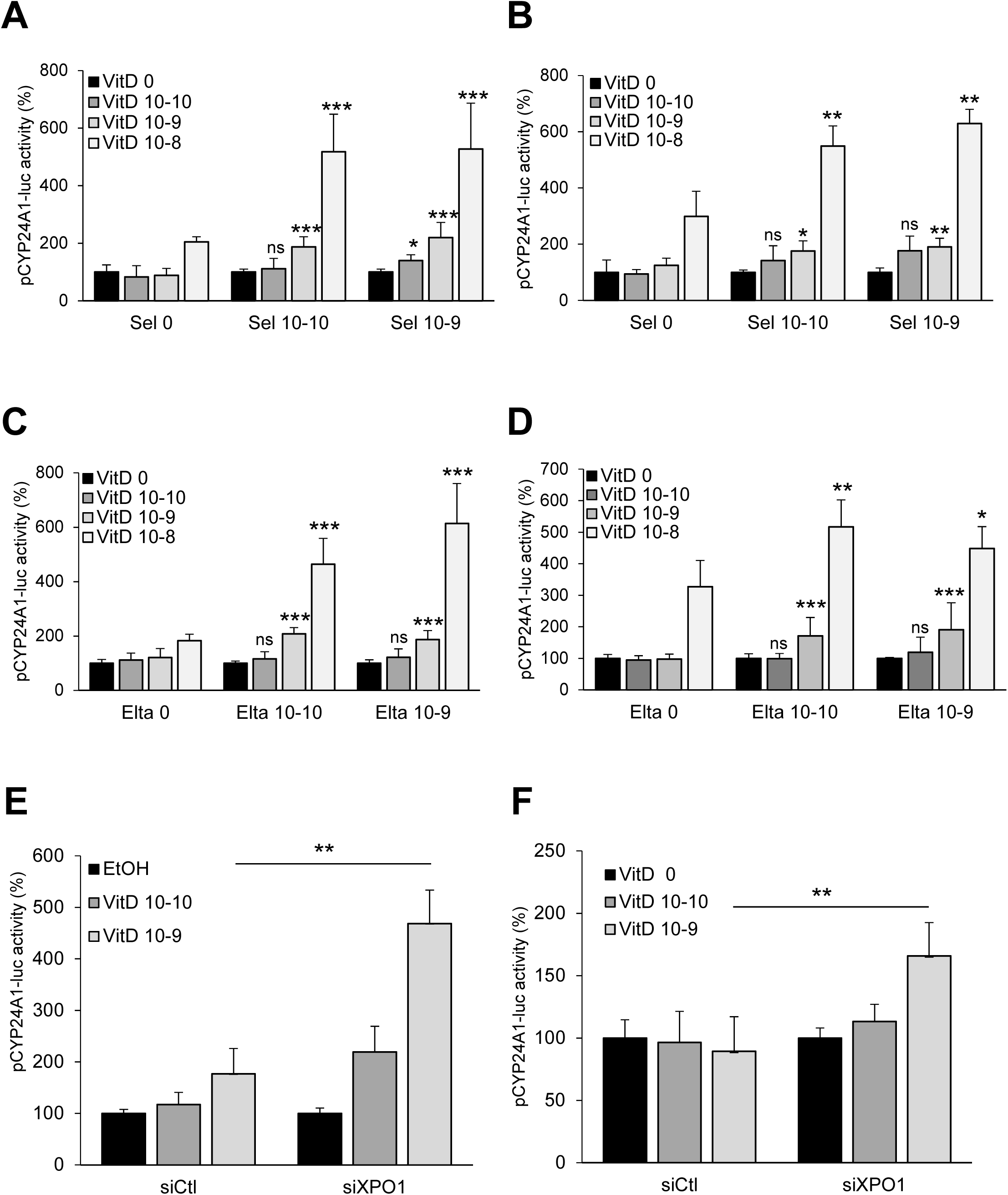
Both Sine and siXPO1 stimulate vitamin-D-dependent activity of CYP24A1 promoter. MDA-MB-436 (A, C and E) and MDA-MB-231 (B, D and F) were transfected with pRL-CMV-renilla and pCYP24A1-luc plasmids and treated with selinexor (A and B) or eltanexor (C and D) and vitamin D or transfected with siXPO1 and treated with vitamin D (E and F). Luciferase values were normalized to the renilla luciferase control and expressed relatively to VitD 0/Sel 0 or Elta 0 or siCtl/EtOH conditions. Statistical analyses were performed using the Mann-Whitney test. *p<0.05, **p<0.01; ***p<0.001.

### Targeting XPO1 increases vitamin D-dependent transcriptional activity of VDR

We next investigated whether XPO1 modulates VDR transcriptional activity. We first used the pCYP24A1-luc reporter, which contains the promoter of the vitamin D–responsive gene CYP24A1 upstream of the luciferase-encoding gene. MDA-MB-436 (Figure 4A and C) and MDA-MB-231 (Figure 4B and D) cells were transfected with the reporter and treated with SINE compounds and vitamin D. Selinexor (10^-10^ and 10^-9^ M) significantly enhanced vitamin D–induced reporter activity in both cell lines (Figure 4A and B). Similar results were obtained with eltanexor (Figure 4C and D), indicating that inhibition of XPO1 activity enhances VDR transcriptional activity.

**Figure 4.**
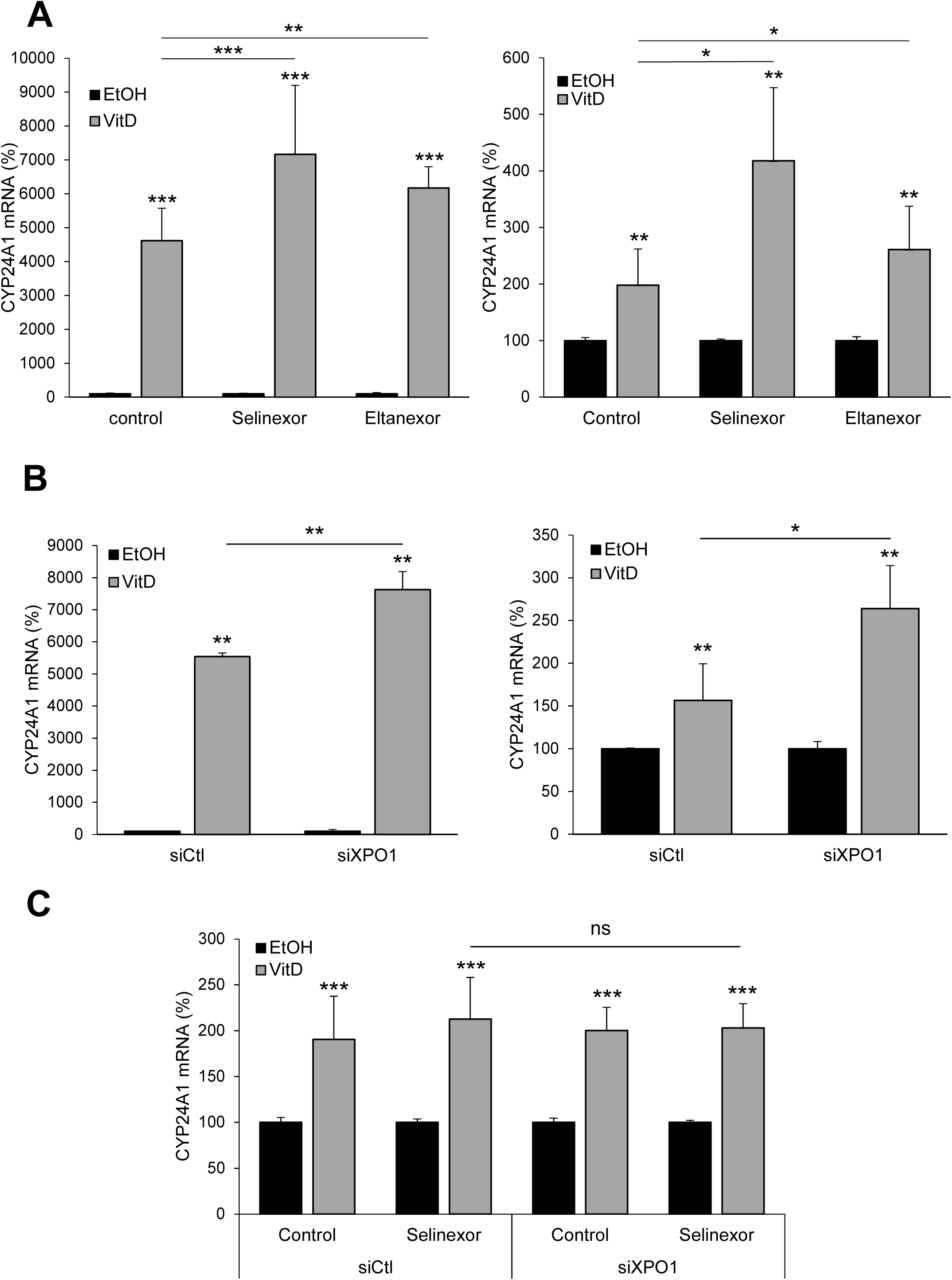
SINEs or siXPO1 augment vitamin D-dependent expression of CYP24A1 mRNA. MDA-MB-436 (left panels) and MDA-MB-231 (right panels) were - treated with either selinexor (10^-8^ M) or eltanexor (10^-8^ M) and/or vitamin D (10^-9^ M) (A) or transfected with siXPO1 and then treated with vitamin D (10^-9^ M) (B). (C) MDA-MB-436 cells were transfected or not with siXPO1 and then treated with selinexor (10^-8^ M) and/or vitamin D (10^-9^ M). After 24h treatment, RNAs were extracted and subjected to RT-qPCR assays, and quantification of CYP24A1 mRNA was expressed relatively to control/ethanol-treated condition. Statistical analyses were performed using the Mann-Whitney test. *p<0.05, **p<0.01; ***p<0.001.

Consistently, XPO1 knockdown increased vitamin D–induced reporter activity, with a stronger effect observed in MDA-MB-436 (Figure 4E) than in MDA-MB-231 cells (Figure 4F). However, combining siXPO1 with selinexor did not produce additional or synergistic effects (Supplementary Figure 2A), suggesting overlapping mechanisms. Similar results were obtained using an independent siRNA (siXPO1#2) (Supplementary Figure 2B).

We then assessed endogenous VDR target gene expression by measuring CYP24A1 mRNA levels. Vitamin D induced CYP24A1 expression more strongly in MDA-MB-436 (46-fold) than in MDA-MB-231 cells (2-fold) (Figure 5A). In both cell lines, SINE treatment further enhanced this induction. Similarly, XPO1 knockdown increased vitamin D–dependent CYP24A1 expression (Figure 5B), but combining siXPO1 with selinexor did not yield additional effects (Figure 5C). These findings were confirmed using siXPO1#2 (Supplementary Figure 3 A and B) and extended to another TNBC cell line MDA-MB-468 and a luminal breast cancer cell line MCF7 cells (Supplementary Figure 3C and D). Altogether, these data indicate that XPO1 negatively regulates vitamin D–dependent VDR transcriptional activity.

**Figure 5.**
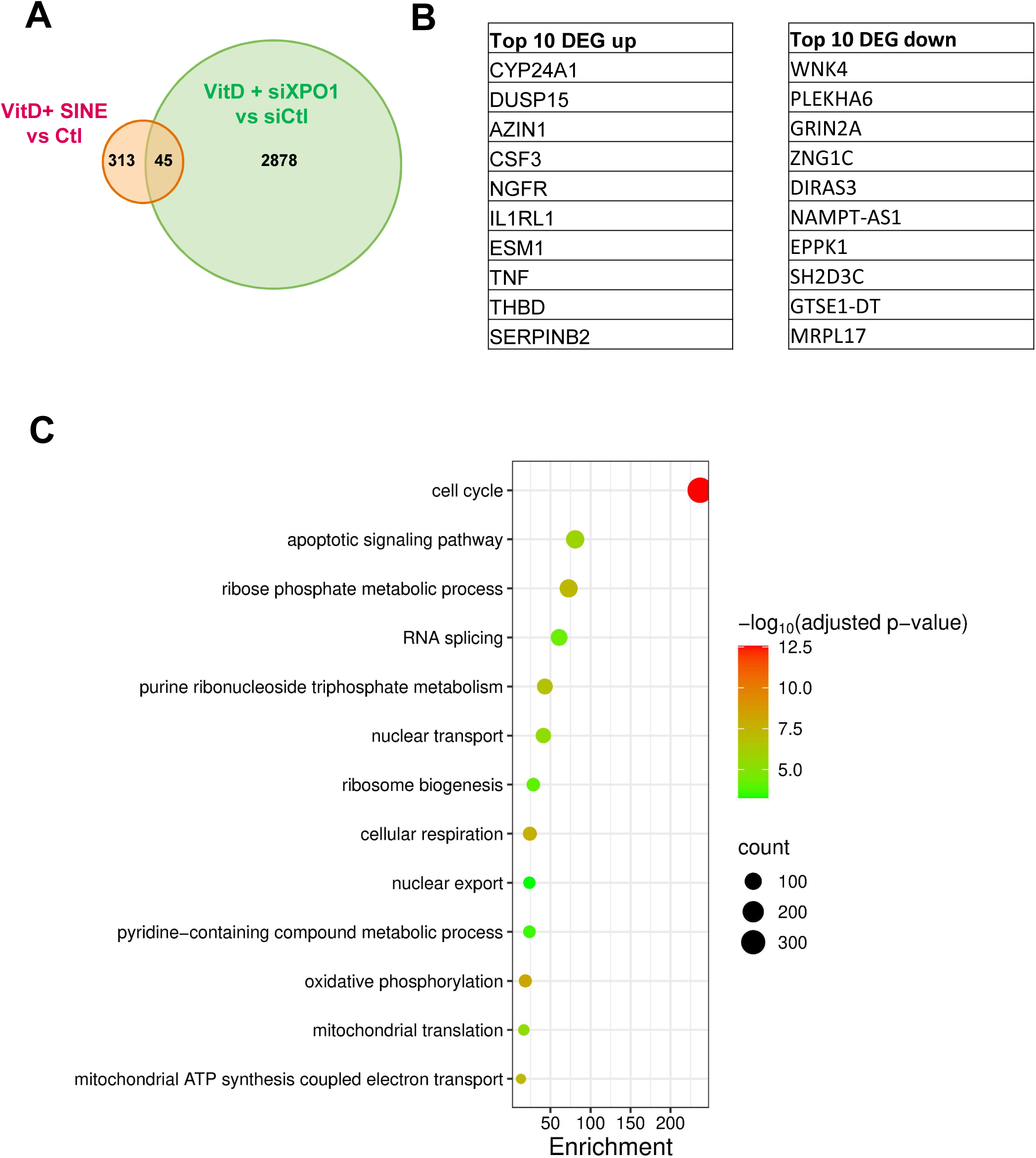
Identification of genes regulated by the combination vitamin D/downregulation of XPO1 in MDA-MB-436 cells. (A) Venn diagram showing the non-redundant set of genes the combination of vitamin D treatment and XPO1 knockdown. Differentially expressed genes (DEGs) were identified from the following comparisons: vitamin D (VitD) + selinexor (Sel) vs control (Ctl) and VitD + eltanexor (Elta) vs Ctl are represented as VitD + SINE vs Ctl, and siXPO1 vs siCtl. DEG lists were combined to remove duplicates. (B) Left panel: Top 10 upregulated genes identified among the DEG sets represented in panel A. Right panel: Top 10 downregulated genes identified among the DEG sets represented in panel A. (C) Gene Ontology Biological Process enrichment analysis of the DEGs identified in panel A. Dot size represents the number of genes associated with each GO term, the x-axis indicates the enrichment score, and dot color represents statistical significance as −log 10(adjusted p-value).

**Figure 6.**
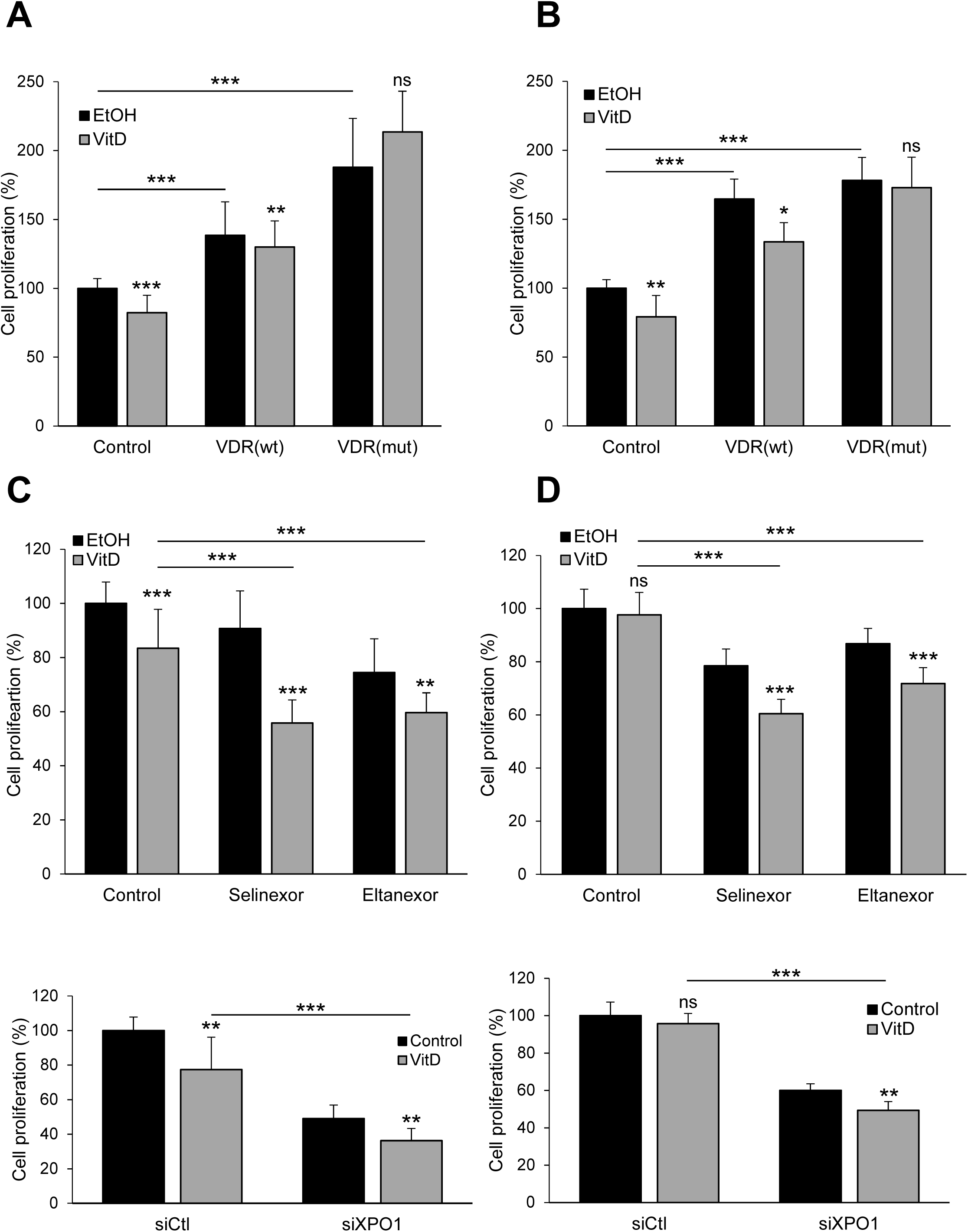
Impaired XPO1 increases vitamin D-dependent repression of TNBC cell proliferation. MDA-MB-436 (left panels) and MDA-MB-231 (right panels) were transfected with either wild type or mutated VDR expression vectors (A). The cells were treated with either selinexor (10^-8^ M) or eltanexor (10^-8^ M) and/or vitamin D (10^-9^ M) (B) or transfected with siXPO1 and then treated with vitamin D (10^-9^ M) (C). Cell proliferation was measured by resazurin reduction in metabolically active cells. Fluorescence (560-590 nm) intensity, proportional to viable cell number, was quantified after one hour of incubation with the dye. Results were expressed relatively to control/ethanol-treated condition. Statistical analyses were performed using the Mann-Whitney test. *p<0.05, **p<0.01; ***p<0.001.

Since vitamin D was shown to repress epithelial-to-mesenchymal transition (EMT)^20^, we investigated whether a combination of vitamin D and XPO1 inhibitors could regulate the expression of CDH1 gene. As shown in Supplementary Figure 3E and F, the combination of SINE compounds or siXPO1 with vitamin D significantly increased the expression of the epithelial marker *CDH1*.

### RNA sequencing analysis of genes responding to the combination of vitamin D treatment and XPO1 knockdown

To obtain a comprehensive view of the genes regulated by the combination of vitamin D treatment and downregulation of XPO1 activity or expression, we performed RNA sequencing. To this end, MDA-MB-436 cells were treated with vitamin D in combination with SINE (selinexor or eltanexor), or with siXPO1.

As shown in Figure 5A, the number of genes differentially regulated by the combination vitamin D/downregulation of XPO1 expression (2923 genes) was substantially higher than that observed with vitamin D in combination with selinexor or eltanexor, collectively referred to as vitamin D/SINE treatment (358 genes). Moreover, only 45 genes were commonly regulated by vitamin D/siXPO1 and vitamin D/SINE. Interestingly, among the differentially regulated genes, *CYP24A1* emerged as the most strongly up-regulated gene (Figure 5B). As mentioned above, CYP24A1 is a *bona fide* VDR responsive gene ^21^ which limits the amount of circulating vitamin D by catalyzing the conversion of 1,25(OH)2D3 into 24-hydroxylated products.

Gene ontology analysis (Figure 5C) of the 3281 genes presented in Figure 5A revealed numerous cellular pathways affected by the combination vitamin D treatment and downregulation of XPO1. Among these, cell cycle and apoptotic signaling pathway were the two main categories. Vitamin D has long been known as a regulator of the cell cycle, notably by stimulating the expression of the cyclin-dependent kinase inhibitors p21/CDKN1A and p27/CDKN1B ^22^. Vitamin D has also been described to induce biochemical alterations associated with apoptosis, including DNA breakage and chromatin condensation, generation of reactive oxygen species and mitochondria disruption ^23^.

### XPO1 regulates vitamin D-driven repression of cell proliferation

Cell cycle regulation coming in the top pathways regulated, we questioned the effect of these combinations on proliferation. First, we examined whether cytoplasmic VDR itself could regulate this process. MDA-MB-436 and MDA-MB-231 cells were transfected with either wild-type VDR (wtVDR) or a nuclear localization signal–deficient mutant (VDR(mut)) ^24^, which is unable to translocate to the nucleus, and then treated with vitamin D (Figure 7A and B). In cells transfected with the empty vector (Control), vitamin D reduced proliferation in both cell lines. Overexpression of VDR(wt) increased proliferation in the absence of ligand, consistent with previous observations in MCF7 cells ^25^, while vitamin D treatment still inhibited this effect. In the absence of ligand, expression of VDR(mut) led to a stronger increase in proliferation in MDA-MB-436 cells and abolished the anti-proliferative effect of vitamin D in both TNBC cell lines. These results indicate that cytoplasmic VDR promotes TNBC cell proliferation, in agreement with our hypothesis.

**Figure 7.**
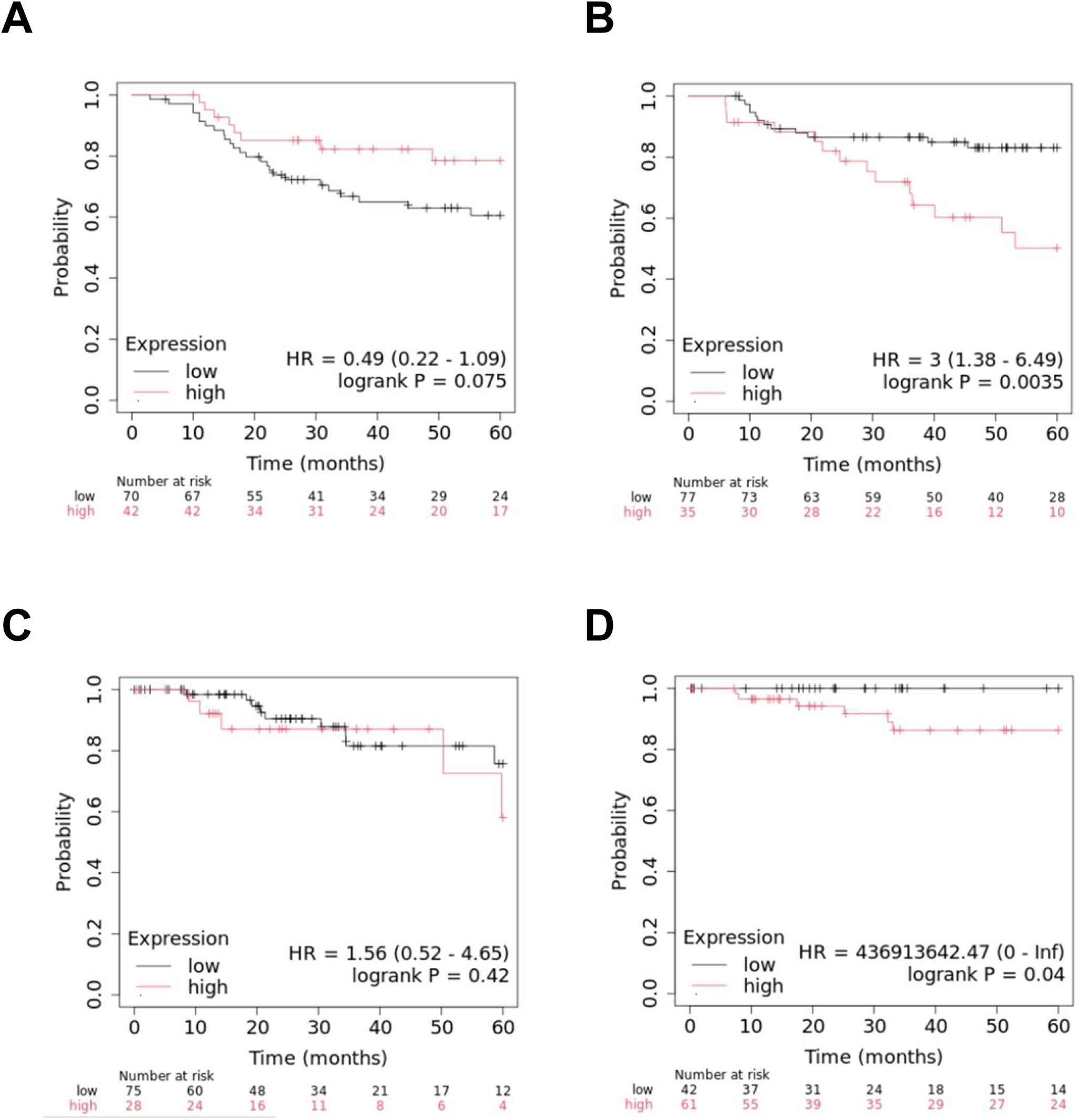
XPO1 expression is associated with good prognosis only when VDR is low. The Kaplan-Meier method was used to estimate overall survival of patients from the RFS Affymetrix (A and B) and the TCGA dataset (C and D). Patients were classified as expressing low levels (A and C) and high levels (B and C) of XPO1./

We then questioned whether XPO1 inhibition affects vitamin D–dependent proliferation. In both MDA-MB-436 and MDA-MB-231 cells (Figure 7C and D), treatment with selinexor or eltanexor enhanced the anti-proliferative effect of vitamin D. Similarly, XPO1 knockdown (Figure 7E and F) increased the ability of vitamin D to suppress cell proliferation. These findings demonstrate that XPO1 is a key regulator of VDR-dependent TNBC cell proliferation.

### XPO1 expression influences the prognosis value of VDR in TNBC

To validate the above-mentioned results using patient samples, we asked whether XPO1 expression could influence the prognostic value of VDR in TNBC patients. We used KM-plotter database and two transcriptomic datasets. As shown in Figure 1A and B, using Affymetrix dataset, we observed that high VDR expression was significantly associated with short recurrence-free survival (RFS) of patients with tumors exhibiting high XPO1 mRNA expression (panel B). In patients, with low XPO1 mRNA expression (panel A), VDR expression was rather correlated with a better RFS although the correlation not significant. Similar results were obtained in a second dataset obtained by RNA sequencing and by analyzing patient overall survival (Figure 1 C and D). These data supported the fact that XPO1 indeed control VDR biological activity by regulating its subcellular localization as supported by the present work.

## DISCUSSION

In this study, we demonstrated that relocalization of VDR in TNBC cells by repressing the activity or the expression of XPO1 was able to increase vitamin D-dependent activity of the receptor. We have been able to show that VDR could then induce more strongly the expression of vitamin D responsive gene as well as repress more efficiently TNBC cell proliferation.

Overall VDR expression is generally considered beneficial in breast cancer. A high immunoreactivity score has been shown to be associated with significantly better progression-free survival and overall survival ^26^. A separate study reported that high VDR expression in invasive breast tumors was associated with favorable prognostic factors and a low risk of breast cancer death ^27^. It was recently reported that, in TNBC, high VDR expression was observed in both the cytoplasm and the nucleus, whereas luminal B tumors exhibited high VDR expression exclusively in the cytoplasm ^28^. Moreover, nuclear VDR expression correlated with low-grade tumors, while cytoplasmic expression correlated with lymph node-positive tumors ^29^. In patients with TNBC, cytoplasmic VDR expression showed a significant inverse correlation with disease-free survival ^12^.

The subcellular localization of VDR also influences outcomes in other cancers. Indeed, high cytoplasmic VDR expression in ovarian cancer was associated with impaired overall survival ^30^. It has also been described that nuclear VDR expression is strongest in normal colorectal mucosa and in hyperplastic polyps, whereas it is extremely weak or absent in colorectal carcinoma. At the same time, cytoplasmic VDR is weakest in normal colorectal mucosa and hyperplastic polyps, increases during the adenoma stage, and is most abundant in colorectal carcinoma ^31^.

The observation that cytoplasmic VDR expression was associated with a poor prognosis in patients with TNBC is reminiscent of what we previously described for the two transcription factors RIP140 (receptor-interacting protein 140) and LCoR (ligand-dependent corepressor) ^32^. Nuclear expression of RIP140 and LCoR was negatively correlated with breast tumor size, whereas their cytoplasmic expression was associated with that of tumor aggressiveness markers. In prostate cancer, predominant nuclear expression of AR (androgen receptor) is associated with non-neoplastic tissues, whereas cytoplasmic AR expression was highest in androgen-independent prostate cancer ^33^. The subcellular localization of β-catenin was also linked to colorectal cancer: nuclear expression was never found in normal tissues, whereas it was observed in 100% of carcinomas ^34^. The expression level of cancer markers has long been a paradigm in cancer prognosis, but subcellular localization is receiving increasing attention.

Selinexor was first used in the clinic to treat patients with multiple myeloma, in combination with bortezomib and dexamethasone ^35^. Selinexor has also been evaluated in other hematologic cancers, such as advanced B-cell malignancies ^36^, showing tolerability in patients in phase I trials. It was also tested in phase I trials in patients with TNBC in combination with eribulin ^37^. The combination was well tolerated, with mild to moderate adverse events, and produced antitumor responses. Preclinical studies indicated that selinexor is able to augment eribulin’s cytotoxic and apoptotic effects ^38^.

XPO1 has also been implicated in drug resistance, and combinations of selinexor with various anticancer agents have been tested in solid tumors. Very recently, a study reporteded the use of selinexor in combination with gemcitabine to treat patients with advanced sarcomas ^39^. The phase I study indicated that patients’ overall health status did not deteriorate, and the most frequent adverse events were neutropenia and thrombocytopenia. *In vitro* and *in vivo* models of glioblastoma have shown that selinexor enhances the radiosensitivity of cancer cells ^40^. On this basis, a phase I trial was conducted to evaluate the safety, tolerability, and preliminary efficacy of selinexor combined with chemoradiotherapy in patients with newly diagnosed glioblastoma ^41^, confirming the safety of the treatment. Given the anti-oncogenic properties of vitamin D, vitamin D-based compounds have also been investigated as potential therapeutic agents in various cancers. A phase I trial was conducted in patients with metastatic pancreatic cancer ^42^. Patients received paclitaxel and paricalcitol, a synthetic VDR agonist which proved to be globally safe and induced favorable changes in the tumor microenvironment. Conversely, very recently, in a phase 2 randomized clinical trial, high-dose vitamin D3 added to standard treatment in patients with metastatic colorectal cancer did not improve progression-free survival ^43^.

When we performed RNA sequencing, among the different treatments were selinexor alone and eltanexor alone. Since the two drugs, long-described inhibitors of XPO1 activity, share a similar structure, it was expected that a large proportion of genes would be commonly targeted by the two compounds. Unexpectedly, when used alone, Venn diagram analysis revealed that the overlap between the two conditions was modest, with only 13 genes (data not shown). Eltanexor was originally designed to achieve lower central nervous system exposure and improved gastrointestinal tolerability compared with selinexor, in order to enable more frequent and prolonged dosing ^44^. It can be hypothesized that the clinical differences observed between selinexor and eltanexor are encoded by a large number of genes, further explaining the modest overlap in the Venn diagrams.

Very recently, a highly promising compound has been described as a new reversible XPO1 inhibitor suitable for cancer therapy ^45^. This compound (known as FR-027) targets Cys528 through nucleophilic aromatic substitution. In vivo, in xenograft and syngeneic models, the compound delayed disease progression while increasing overall survival across hematological and solid tumor models. Notably, FR-027 is not associated with adverse secondary effects, likely due to the reversibility of its interaction with XPO1 ^45^. This study justifies further investigation toward the clinical development of FR-027.

## ACKNOWLEDGEMENTS

We would like to thank Pr Ana Aranda for the kind gift of pCYP24A1-luc and Pr Guan Chen for providing the vectors encoding wild-type VDR and the NLS-mutant form of VDR.

## AUTHORS CONTRIBUTIONS

JD and SB investigated. BG, UJ and CT contributed to formal analysis, VC participated to conceptualization, investigation, formal analysis, writing-original draft and funding acquisition and SJ participated to conceptualization, investigation, formal analysis, methodology writing-original draft and funding acquisition.

## DATA AVAILABILITY STATEMENT

The data supporting the findings of our study are available from the corresponding author upon reasonable request.

## Abbreviations

BC: breast cancer
TNBC: triple negative breast cancer
VDR: vitamin D receptor
XPO1: exportin-1.

