## Supplementary figures for "XPO1 controls VDR localization and activity in triple-negative breast cancer cells"

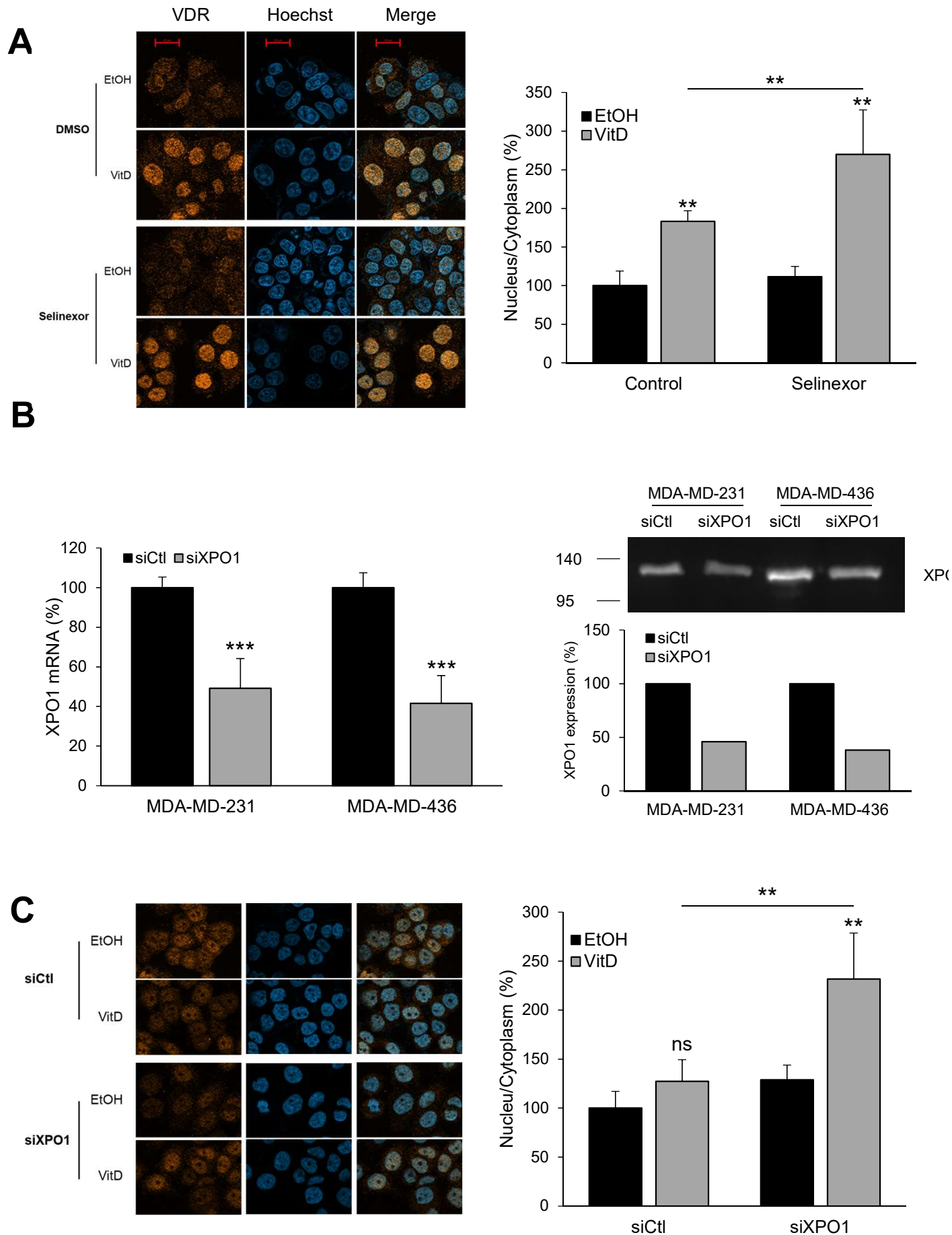

Supplementary Figure 1

**Supplementary Figure 1.** A. Selinexor increases vitamin D-dependent nuclear expression of VDR in MCF7 cells. The cells were treated with selinexor ($10^{-8}$ M) and/or vitamin D ($10^{-9}$ M). Immunofluorescence staining (left panels) was performed with an anti-VDR antibody and nuclei were counterstained using Hoechst. Cells were fixed after 24 h of treatment, and images were acquired at 40× magnification using a Zeiss AxioImager M2 Apotome2. Staining quantification (right panels) was performed using Fiji software, and results are expressed as a percentage relative to the control/ethanol (EtOH)-treated condition. B. MDA-MB-231 and MDA-MB-436 were transfected with either siCtl or siXPO1. Left panel. RNAs were extracted, subjected to RT-qPCR assays and quantification of XPO1 mRNA was expressed relatively to siCtl condition. Right panel. Protein extracts were submitted to western blotting using an anti-XPO1 antibody. C. siXPO1 increases vitamin D-dependent nuclear expression of VDR in MDA-MB-468. The cells were transfected with siXPO1 and then treated with vitamin D ($10^{-9}$ M). Immunofluorescence staining (left panel) and quantification (right panel) were done as in (A). Statistical analyses were performed using the Mann-Whitney test. \*\*$p < 0.01$. \*\*\*$p < 0.01$.

**A**

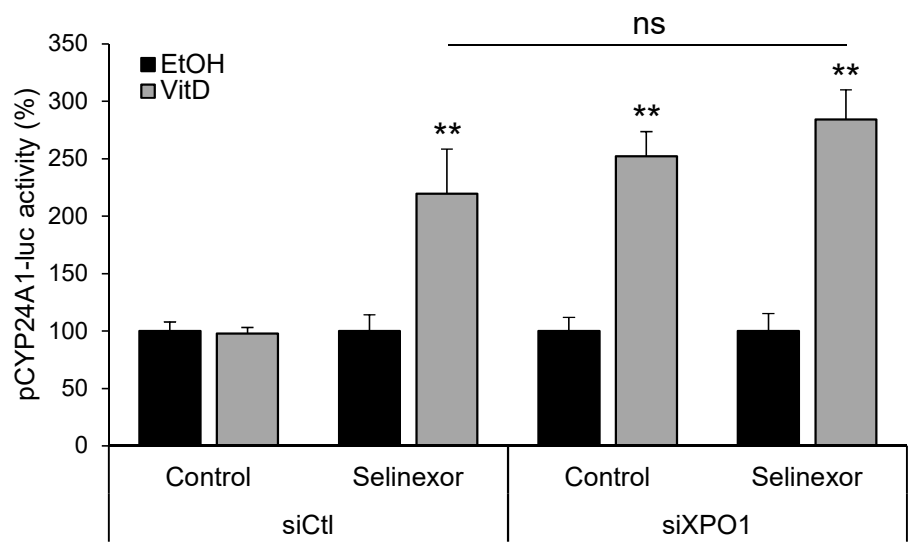

**B**

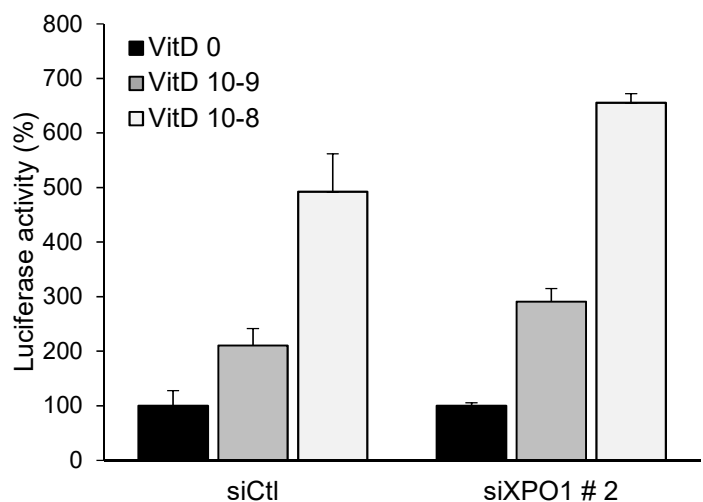

**Supplementary Figure 2.** (A) MDA-MB-436 cells were transfected with pRL-CMV-renilla and pCYP24A1-luc plasmids and siXPO1 and then treated or not with selinexor and/or vitamin D. Luciferase values were normalized to the renilla luciferase control and expressed relatively to the siCtl/ethanol (EtOH)-treated condition. (B) MDA-MB-231 cells were transfected with either siCtl or siXPO1#2 as well as pRL-CMV-renilla and pCYP24A1-luc plasmids and then treated with vitamin D. Luciferase values were expressed as in (A). Statistical analyses were performed using the Mann-Whitney test. \*\*p<0.01.

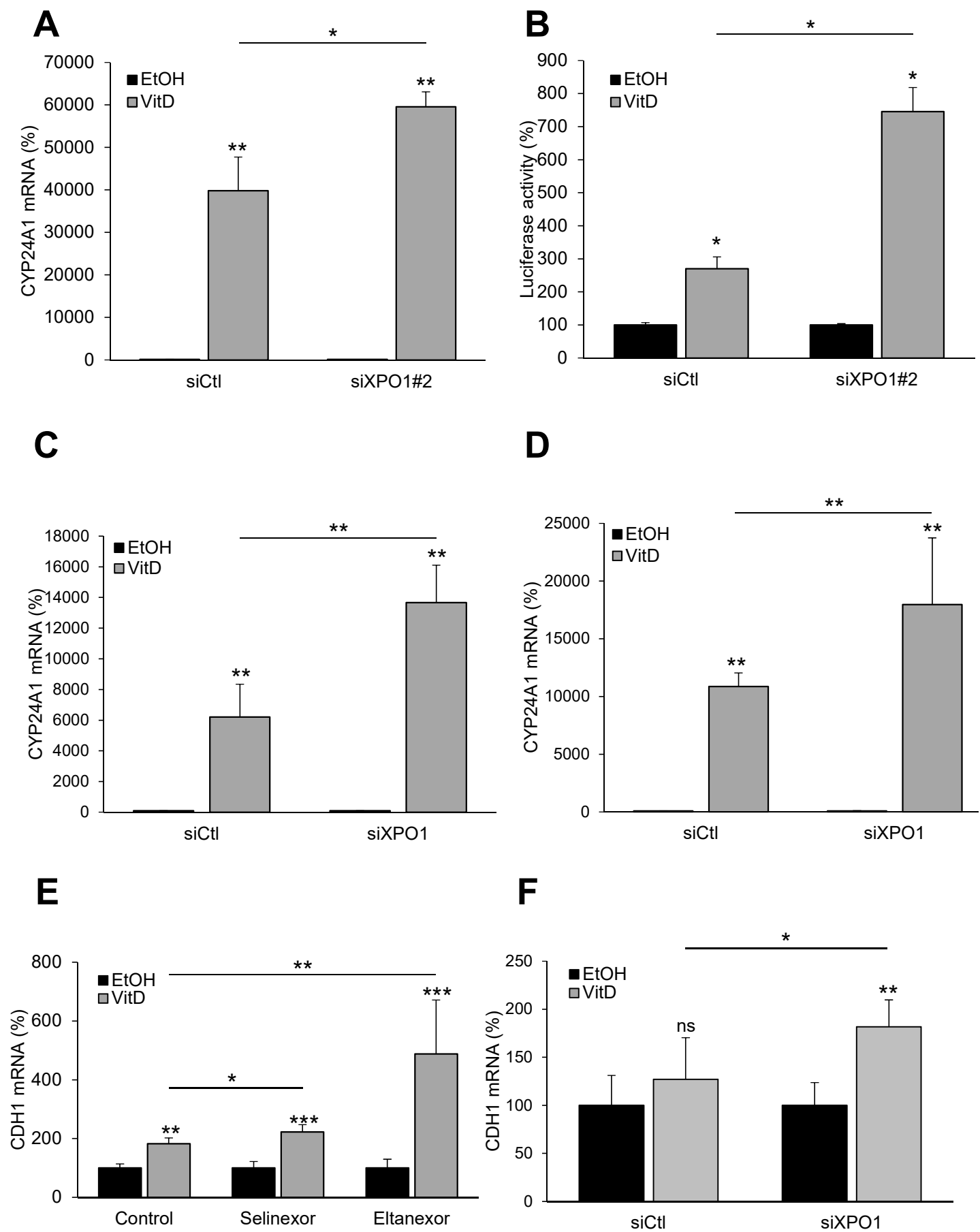

**Supplementary Figure 3**

**Supplementary Figure 3.** MDA-MB-436 (A) and MDA-MB-231 (B) cells were transfected with either siCtl or siXPO1#2 and then treated with vitamin D ($10^{-9}$ M). RNAs were extracted and subjected to RT-qPCR assays, and quantification of CYP24A1 mRNA was expressed relatively to control/ethanol-treated condition. MDA-MB-468 (C) and MCF7 (D) were transfected with either siCtl or siXPO1 and treated or not with vitamin D ($10^{-9}$ M). After 24h treatment, RNAs were extracted and subjected to RT-qPCR assays, and quantification of CYP24A1 mRNA was expressed relatively to siCtl/ethanol-treated condition. MDA-MB-231 were treated with either selinexor ($10^{-8}$ M), eltanexor ($10^{-8}$ M) and/or with vitamin D ($10^{-9}$ M) (E) or transfected with siCtl or siXPO1 and then treated with vitamin D ($10^{-9}$ M) (F). CDH1 quantification was performed as explained above. Statistical analyses were performed using the Mann-Whitney test. \*\*$p < 0.01$. \*\*$p < 0.01$. \*\*\*$p < 0.01$.
